# Catching more air: A method to spatially quantify aerial triazole resistance in *Aspergillus fumigatus*

**DOI:** 10.1101/2022.11.03.515058

**Authors:** Hylke H. Kortenbosch, Fabienne van Leuven, Cathy van den Heuvel, Sijmen S. Schoustra, Bas J. Zwaan, Eveline Snelders

## Abstract

Airborne triazole-resistant spores of the human fungal pathogen *Aspergillus fumigatus* are a significant human health problem as the agricultural use of triazoles has selected for cross-resistance to life-saving clinical triazoles. However, how to measure the health risk posed by these inhaled spores remains unclear. Here, we describe a method for cost-effective wide-scale outdoor air sampling to measure both spore abundance as well as antifungal resistance fractions. We show that prolonged outdoor exposure of sticky seals placed in delta traps, when combined with a two-layered cultivation approach, can consistently yield sufficient colony-forming units (CFUs) for the quantitative assessment of aerial resistance levels at a spatial scale that was up to now unfeasible. When testing our method in a European pilot sampling of 12 regions, we demonstrate that the triazole-resistant fraction of airborne spores is widespread and varies between 0 and 0.1 for itraconazole (*∼*4 mg/L) and voriconazole (*∼*2 mg/L). This method facilitates the assessment of health risks by pinpointing potential hot- and coldspots of environmental resistance selection. This efficient and accessible air sampling protocol opens up extensive options for fine-spatial sampling and surveillance studies.

**IMPORTANCE:** *Aspergillus fumigatus* is an opportunistic fungal pathogen which humans and other animals are primarily exposed to through inhalation. Due to the limited availability of antifungals, resistance to the first choice class of antifungals, the triazoles, in *A. fumigatus* can make infections by this fungus untreatable and uncurable. Here we describe and validate a method that allows for the quantification of airborne resistance fractions and quick genotyping of *A. fumigatus* TR-types. Our pilot study provides proof of concept of the suitability of the method for use by citizen scientists for large-scale spatial air sampling. Spatial air sampling opens up extensive options for surveillance, health-risk assessment, and studying the ecology of *A. fumigatus* and the development of triazole resistance.

## INTRODUCTION

*Aspergillus fumigatus* is a saprophytic fungus that is ubiquitous in soils rich in organic matter [1]. *A. fumigatus* is especially abundant in dead plant material such as plant waste heaps, where its high thermotolerance allows it to grow under the high temperatures generated during the composting process and form vast numbers of asexual conidiospores [2, 3, 4]. Due to their small size and hydrophobic nature, these spores readily disperse through the air when plant waste heaps are disturbed [5, 6]. The plant waste heaps are considered the optimal habitat for *A. fumigatus*, while this fungus also is an opportunistic pathogen that can infect or colonise the lungs of birds, humans and other mammals [7, 1, 8]. Humans are believed to inhale roughly 100 spores per day on average [9, 1], and although the immune systems of most humans efficiently remove these spores, they can cause symptoms ranging from mild allergies to often-lethal acute invasive aspergillosis infections, depending on the health status of an individual’s lungs and their immune system status [10, 1]. Unfortunately, only a limited pool of antifungals is available to treat *A. fumigatus* infections, with triazoles being the most widely used class.

Quantifying growth and antifungal resistance fractions in plant waste material is useful for the identification of sources of environmental resistance [11], but the direct health risk is posed by inhaled airborne spores. Efforts have been made to assess aerial resistance, predominantly in hospital environments [12, 13, 14]. The methods range from the air compaction of 250 L of air on agar plates [12], collection by homemade dust vanes [14] and the overnight exposure of agar plates [13]. All of these methods are useful for the qualitative detection of *A. fumigatus* conidiospores and resistance genotypes and have even been used to quantitatively estimate colony forming units (CFU) per m^3^ [13]. However, the CFUs reported per sample are generally quite low (<10) in these hospital building studies, probably due to HEPA filtration of the air. When estimating a proportion of a natural population, in our case the fraction of triazole-resistant CFUs, an adequate sample size is key for accuracy. Specifically because at low sample sizes, sampling error increases rapidly to the point where the sample estimates become completely uninformative on quantitative differences in proportions, such as for antifungal resistance fractions [15, 16]). However, quantitative assessment of aerial resistance fractions is key for identifying hotspots and coldspots of triazole resistance, possible transmission routes to patients, and the health risks they pose in different geographical regions. This leads to the question; how can we increase *A. fumigatus* CFUs in air samples to accurately estimate triazole resistance fractions?

A limitation of air compaction or other types of active air sampling devices capable of sampling large volumes of air is that they are costly [17]. This is not a hurdle in longitudinal studies at set locations such as hospitals, however, this makes these devices impractical for more extensive environmental surveys that aim to estimate resistance fractions at various geographic scales. Shelton et al. 2023 [18] therefore proposed a sampling method that at low cost collects airborne fungal spores and also at a wide geographical range by involving citizen-scientists. In short, this method involves attaching two halves of a MicroAmp^TM^ clear adhesive film (Applied Biosystems^TM^, UK) to a window sill with poster putties and exposing them to outside air for six to eight hours on a predetermined sampling day. Subsequently, the film was recovered and returned to the lab by Freepost. In the lab, the film was then pressed onto an agar plate and cultured at 43°C. This method proved effective at capturing *A. fumigatus* spores on a nationwide level, but on average yield only 1.25 *A.fumigatus* CFUs per sample [18]. Thus, the number of recovered CFUs was too low to accurately estimate antifungal resistance fractions per individual sample.

Building on the method of Shelton et al. 2023, we also used sticky seals to capture spores, now with exposure times of four weeks. Compared to agar plates, these seals are less prone to drying out, weathering, or unwanted microbial growth. As prolonged exposure also means potentially increased exposure to other microbes, we tested a culturing method in which we poured Flamingo medium [19] selective for *A. fumigatus*, directly on top of exposed seals in two distinct layers. The first layer allows all *A. fumigatus* spores to germinate and grow, while the second layer is selective for resistance against the triazole compound of choice. In addition to developing this culturing approach, we tested it in an international pilot study, including 12 sampled regions, for which we asked fellow scientists to follow a provided protocol supported with instructions via a video. With this, we assessed whether the use of foldable delta traps, protecting the seals from UV light and rain, could be combined with the use of sticky seals in a flexible, accessible and effective protocol for affordable and large-scale outdoor spore capture.

## RESULTS AND DISCUSSION

### Validation layer culturing approach

We tested whether culturing from seals inoculated with *A. fumigatus* conidia by adding Flamingo media with different triazole treatments directly on top would, relative to plating conidia on the agar surface, affect the assay’s utility to visually screen for triazole-resistant *A. fumigatus* colonies. We found that an interaction between the seals and itraconazole strongly diminished suppression of itraconazole-sensitive strains, compared to using itraconazole in Flamingo agar plates without seals. (see Supplement - Effect of seals on triazole selectivity). To address this technical issue, we validated a layered culturing approach where a triazole-negative layer of medium acts as a buffer between the triazole-containing layer and the seal. For the two clinical triazoles itraconazole and voriconazole, we found that culturing with the described two-layer culturing method only allowed colonies of *A. fumigatus* strains with a MIC_95_ *≥*2 mg/L to breach the agar surface of the upper azole-containing layer. The rate at which different *A. fumigatus* strains did so by the end of incubation seemed largely tied to their MIC_95_ value (Fig. 1). Similarly, we tested whether this two-layer culturing approach could also be used to detect resistance to the agricultural triazoles difenoconazole (4 mg/L) and tebuconazole (6 mg/L). However, these triazoles were found to be insufficiently selective when added to the triazole-containing layer during the layered culturing approach (see Tables S1 and S2). We also conducted limited tests with the agricultural triazoles bromuconazole, epoxiconazole, and propiconazole (data not shown), and these compounds also appeared to be ineffective for resistance screening with the layered culturing method.

**FIG 1.**
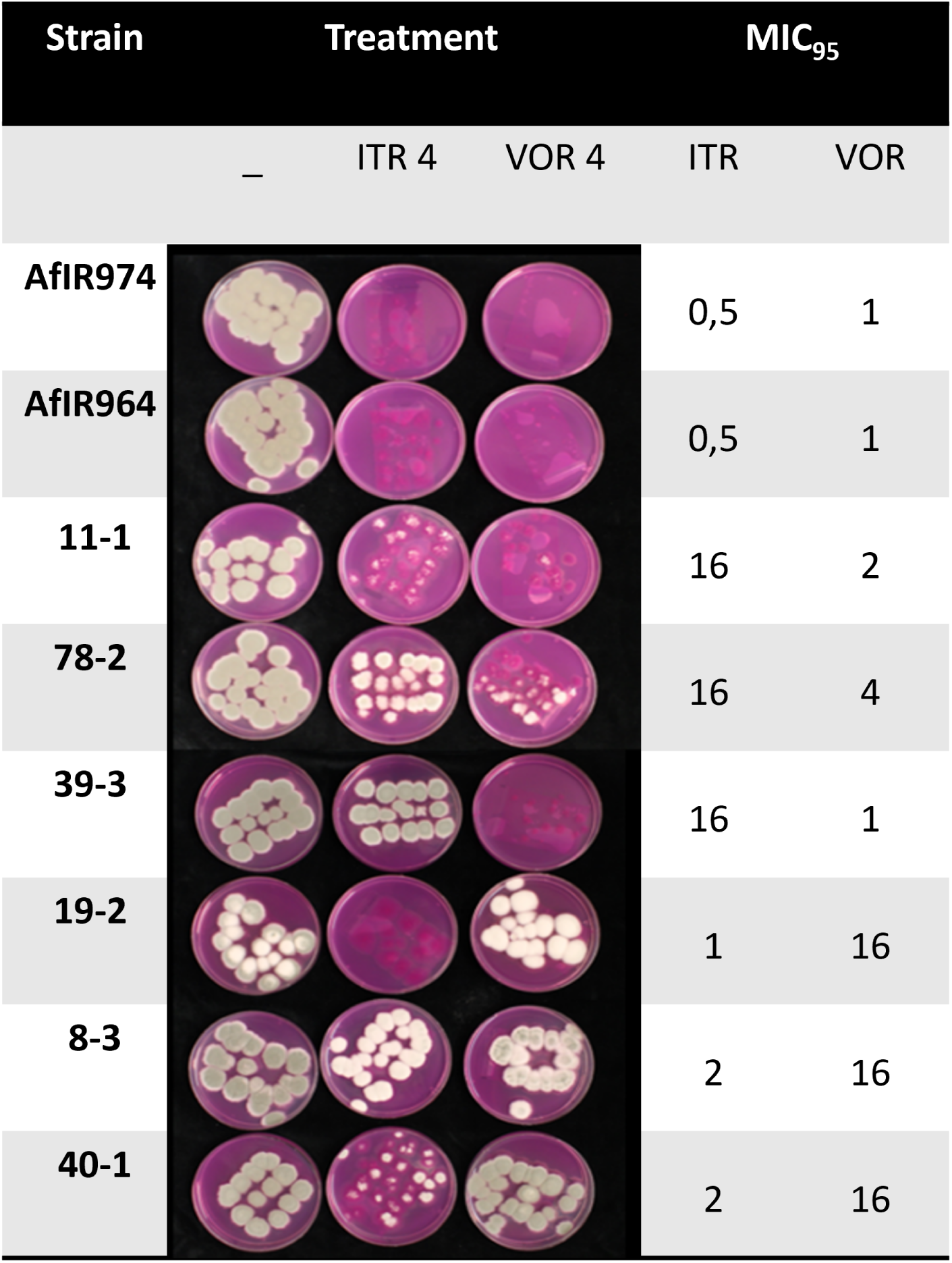
Growth from seals innoculated with *A. fumigatus* conidia of strains with known minimal inhibitory concentration (MIC) values through the top layer of Flamingo medium after eight days of incubation at 48 °C. The included treatments in the upper agar layer are triazole-negative, 4 m/L itraconazole (ITR 4), and 4 mg/L voriconazole (VOR 4). MIC_95_ values in mg/L for itraconazole (ITR) and voriconazole (VOR) of the strains are given in the two right-most columns.

We do not have a full explanation for why some triazoles are applicable in this culturing method and not others. Table 1 lists key characteristics for several medical and agricultural triazole compounds, indicating whether they were tested to be effective with the described layered culturing method to discriminate between azole-susceptible and azole-resistant *A. fumigatus*. Ideally, the applied triazole concentration in the upper agar layer should be higher than the mean MIC_95_ of wild-type strains for discrete selection for resistant strains. The higher mean wild-type MIC_95_ values in agricultural fungicides(Table 1) indicate that they should be present in higher concentrations in the agar to be effective at screening for resistance. Unfortunately, technical aspects of layered culturing directly from sticky seals can make achieving such concentrations difficult. We hypothesize that low solubility in water (Table 1) may cause some triazoles to dissolve in the adhesive of the seal preferentially when applied directly onto the seal in water-based agar, lowering the effective concentration in the agar and reducing the effectiveness of the assay. In the case of itraconazole, its low solubility in water limits its diffusion into the lower triazole-negative layer of the medium. We believe the lower agar layer functions as a buffer here, limiting interaction with the seal. However, in the case of difenoconazole, tebuconazole, and bromuconazole, their intermediate solubilities in water may render this buffering layer less effective. These triazoles may still diffuse into the triazole-negative layer and subsequently bind to the seal, allowing sensitive colonies to grow uninhibited. Voriconazole has a much higher water solubility than itraconazole (Table 1) and likely diffuses into the triazole-negative layer but has seemingly no interaction with the seal (see supplement - Effect of seals on triazole selectivity).

**TABLE 1.**
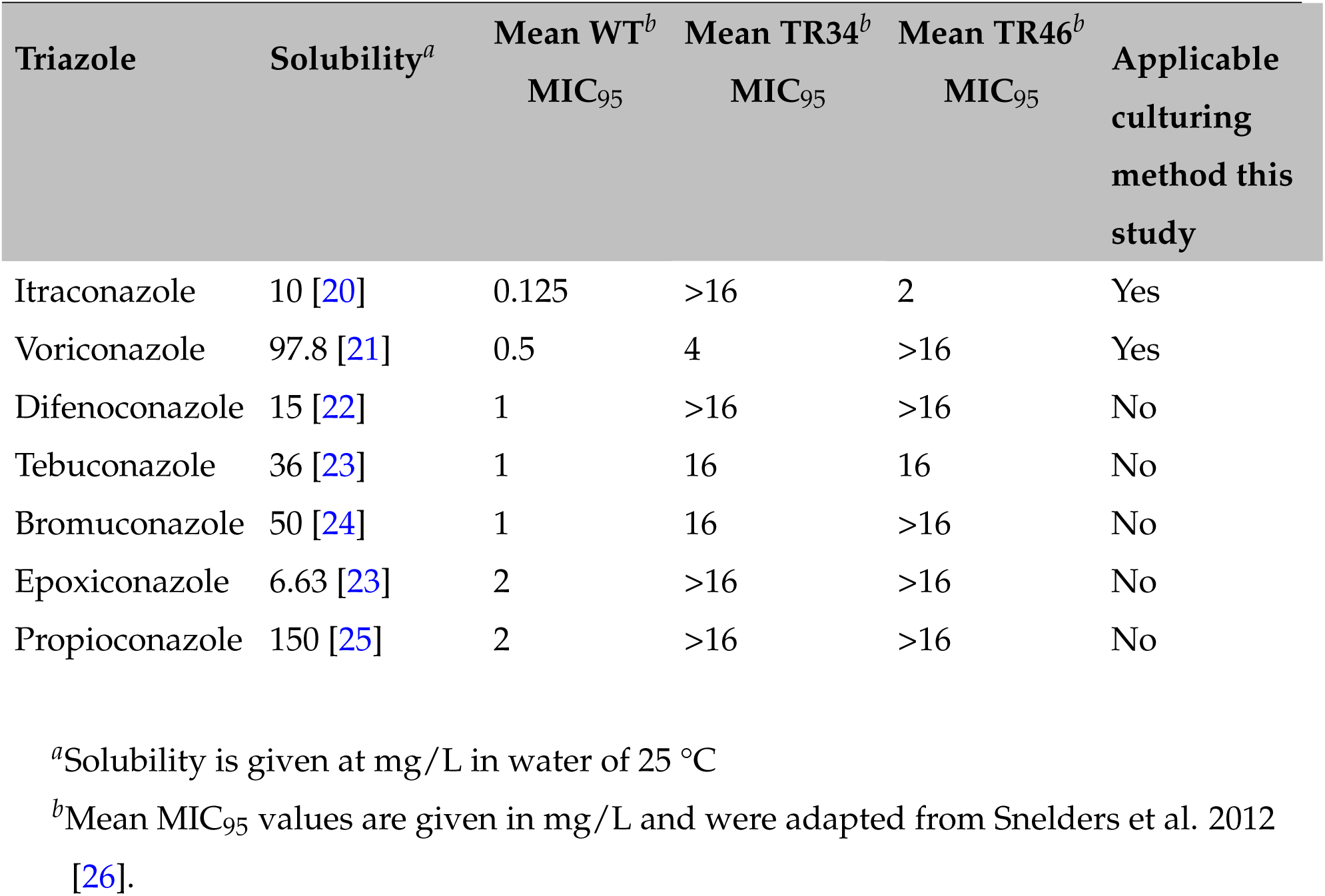
Water solubilities of different triazoles, the mean MIC_95_ values*^a^* for *A. fumigatus* strains with different *cyp*51A genotypes (wild type, TR34/L98H, TR46/Y121F/T289A), and whether the double-layer culturing approach can be used to screen for resistance to these triazoles.

Generally, MIC tests are intended to predict the clinical outcome of treatment of antifungal drugs on isolates under standardised conditions based on established breakpoints [27]. For our purpose of large-scale resistance screening of environmental air samples, we apply the triazoles in Flamingo medium, at 48 ° C, in a non-homogenous (layered) agar, to already germinated colonies growing on the same plate. These culturing conditions are all linked to factors which are known to impact MIC values [27, 28, 29, 30]. The fact that there are inherently no established clinical breakpoints for agricultural fungicides makes finding appropriate concentrations to assay for cross-resistance, and interpreting the output of any such non-standard assay, complex. Therefore, before using this two-layered cultured approach to screen for fungicide resistance it is important to consider the chemical properties of a fungicide. Specifically, its solubility in water, the MIC values of the WT *A. fumigatus* population to it, and its inhibitory effect on *A. fumigatus* when used in the described culturing method.

### International air sampling pilot

Of the 70 air sampling packages that were distributed for the international pilot study, 54 were returned in good order and selected for analysis. These samples were grouped into 12 different regions, in which samples were taken within a circular area with a 50 km radius (Fig. 2). Five of these 54 samples were excluded because of low CFUs (>3x lower than the regional average), three of them were likely taken indoors rather than outdoors. Of the other two samples this was reported to be the case by the participants. Three samples were excluded because of high CFUs (>2x higher than the regional average): one of these traps had fallen from the tree it was hung up in and had been on the ground for an unknown time, and the remaining two were likely deployed close to an *A. fumigatus* growth environment such as a heap of degrading plant waste material. This left 46 representative samples for the analysis of aerial baseline triazole resistance in *A. fumigatus*. To assess regional differences in CFU counts, we also excluded samples deployed later than December 19th 2022 to mitigate the known effect of seasonality on the prevalence of airborne *A. fumigatus*. This left 39 samples for analysis of regional differences.

**FIG 2.**
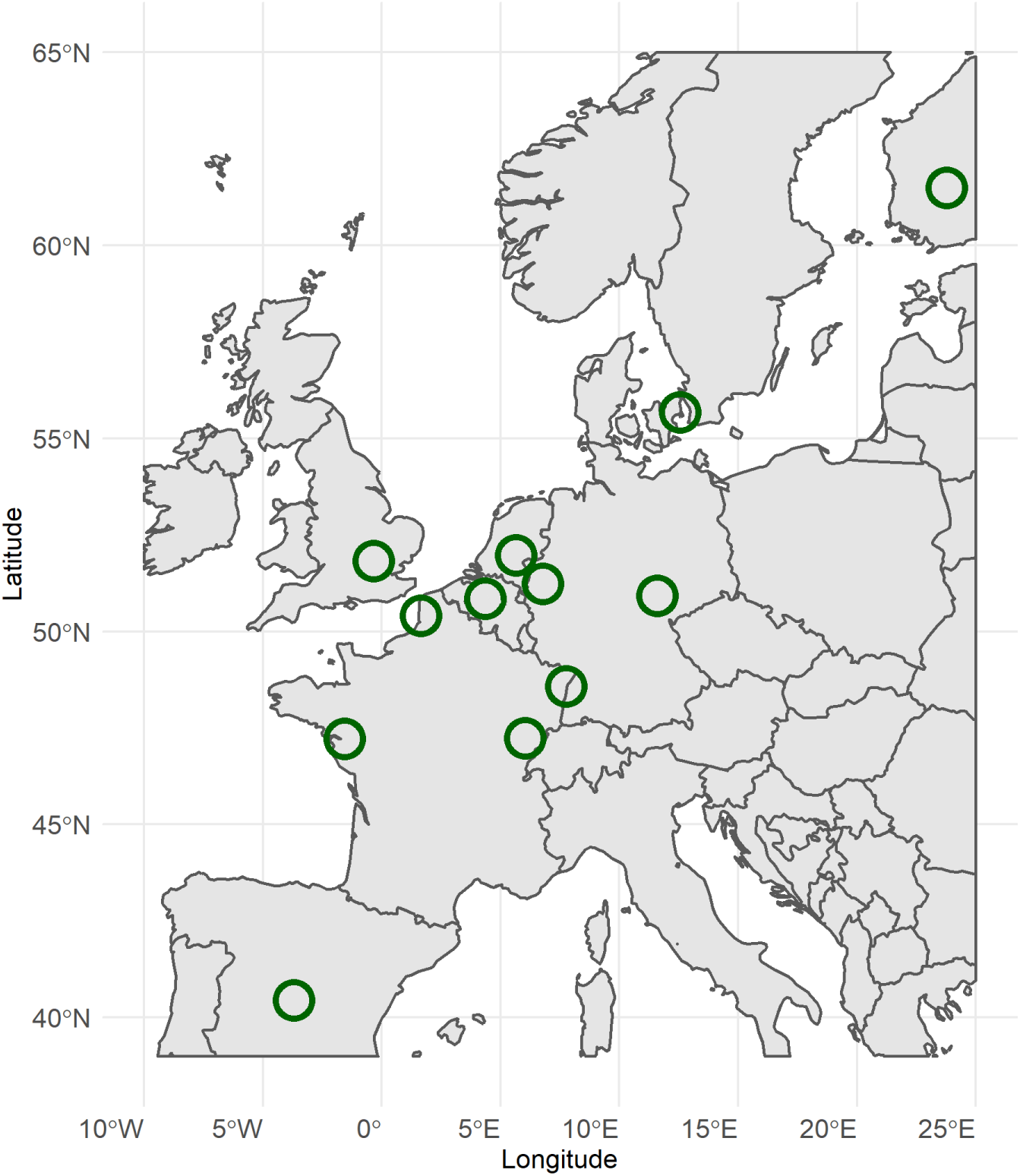
The European regions sampled for the air sampling pilot between November 2022 and January 2023. The green circles designate the regions in which the delta traps were deployed (within a 50 km radius).

### Regional differences in CFU counts

Among the sampled regions, we found significant differences in spore counts between regions (see Fig.3 and Table S3). Samples from The Netherlands, Belgium, and the bordering region of Düsseldorf (Germany) contained 3-fold higher levels of *∼* CFUs than most other regions. We assessed whether weather conditions during the sampling window were significantly correlated with the median CFU counts per trap. We tested for regional averaged UV light, total precipitation, wind speed, and temperature and only found a significant correlation with wind speed (Estimate = 0.5370, SE = 0.1407, z value = 3.818, Pr(>|z|) = 0.000135). This is an expected positive correlation, given that our passive spore capture method requires airflow over the stickers to capture spores. However, in an earlier longitudinal study where more controlled volumes of air were sampled, this correlation was also observed [31]. This suggests that increased CFU counts at greater wind speeds may not simply be a result of more air flowing over the sticker but also reflect an absolute increase in amounts of spores in the air, possibly due to increased disturbance of, and spore release from, *A. fumigatus* sources [1, 32]. Furthermore, the relationship with wind speed is not linear. The regions of Copenhagen, Verton, and Nantes with the highest average wind speed have the lowest counts. These are all coastal regions, which could also have an effect, given that *A. fumigatus* is a terrestrial mould. Moreover, given the limited size of our pilot data and that the regions with the highest counts are adjacent, we cannot rule out spatial auto-correlation in the data. More systematic spatial sampling will be needed under variable weather conditions to separate the factor space from the effect of weather conditions.

**FIG 3.**
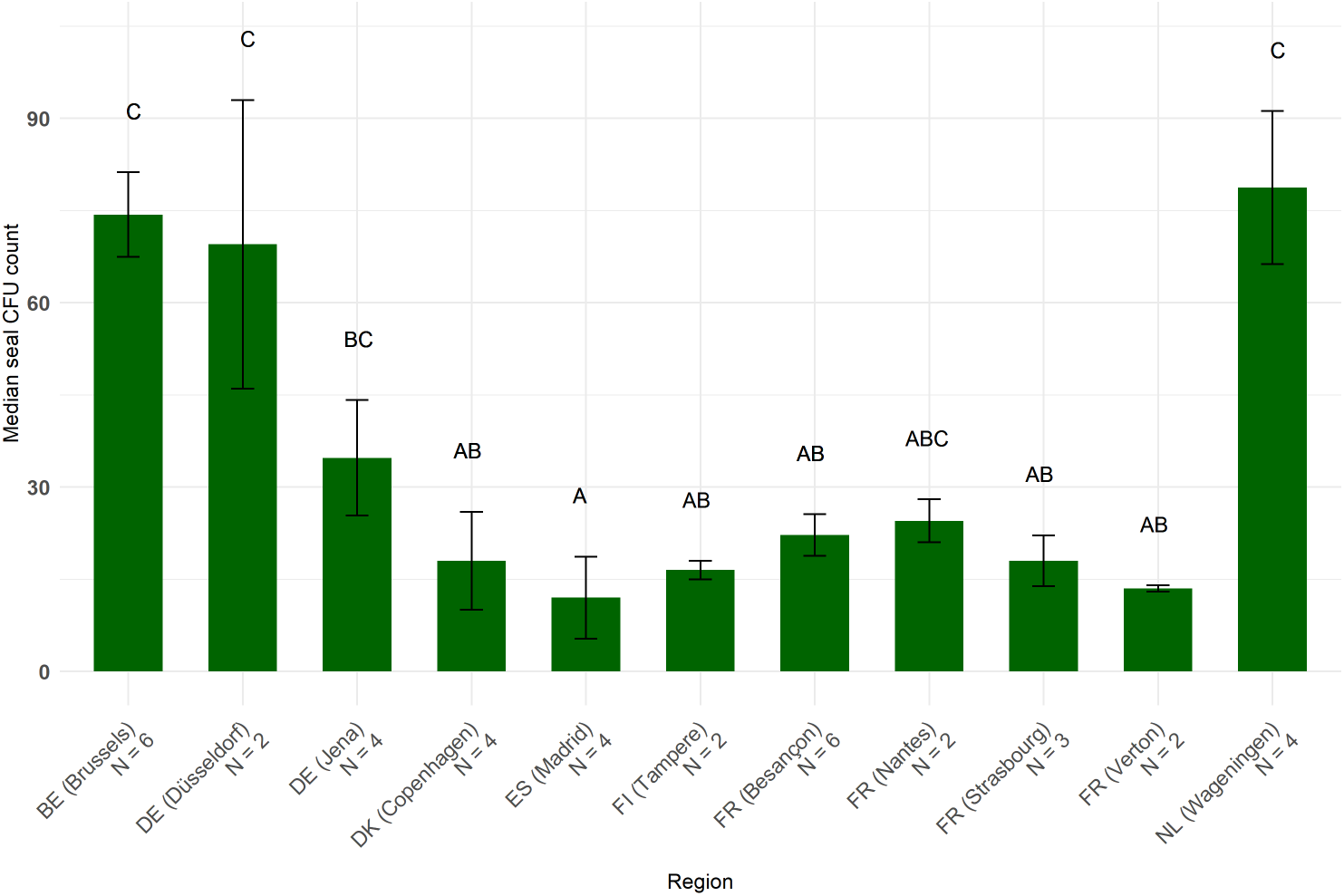
The median CFU count per trap per region (mean *±* SE). The Netherlands, Belgium, and the bordering region of Düsseldorf have noticeably higher CFU counts than most other sampled regions.

Our analysis shows regional differences in CFU capture by the traps. Pilot studies, similar to the one described here should precede systematic sampling for aerial triazole resistance in new regions to assess what number of CFUs can be expected. Due to the passive nature of our air sampling method and the possibility of seasonality in *A. fumigatus* abundance [18], CFU counts may, where necessary be increased by 1) increasing the length of the sampling window and/or 2) adjusting the sampling season. This raises the question: how are these CFU counts per sample relevant for the assessment of resistance fractions?

### How accuracy matters

When estimating a true value in a population, in our case the resistance fraction to triazoles, power calculations are key to determining what number of colonies will provide the desired precision of the population estimate. To determine the number of CFUs (S) needed to estimate a proportion of a population with a binomial distribution the following equation can be used,

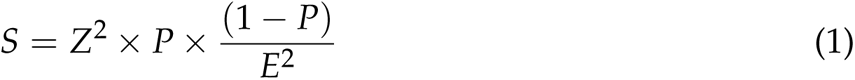

which we can rework to calculate the error size:

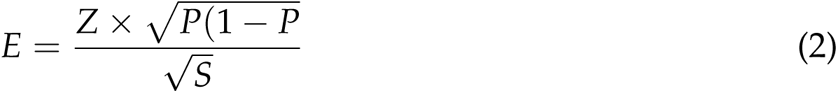

The key factors here are the expected population resistance fraction (P), the sampling error (E) and the critical value (Z) for the desired confidence level of your sample estimate, which is 1.96 for the 95% confidence intervals commonly applied in biology [33]. Based on data from previous studies [34, 35, 18], we expected a resistance fraction of *∼*5%, so will fill in 0.05 as our P. Combining this value with the common 95% confidence interval (Z = 1.96), we can simplify the equation to:

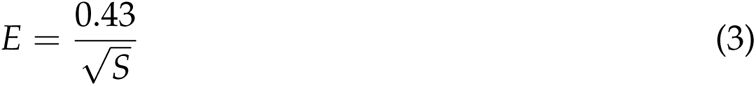

On the one hand, plotting this equation shows that the sampling error, or how much a sample resistance fraction may differ from the true population resistance fraction, rapidly increases at counts <25 (Fig. 4). Previous estimates of baseline resistance based on aerial surveys reported resistance fractions of *∼*0.05 [18, 17]. Therefore, a detection limit of >0.04 is undesirable for quantitative assessment of aerial resistance fractions. Hence, we only included samples that contained *≥*25 CFUs in our quantitative assessment of regional differences in triazole resistance fractions. On the other hand, having over 250 CFUs makes counts less accurate due to overcrowding, and only provides marginal statistical benefits (Fig. 4). This gives us our desired range of 25-250 CFUs per sample. Based on this, only regions with two or more samples in this range were included in our quantitative analysis of resistance fractions. This further restricted our analysis to 22 samples for itraconazole resistance, and 18 for voriconazole resistance.

**FIG 4.**
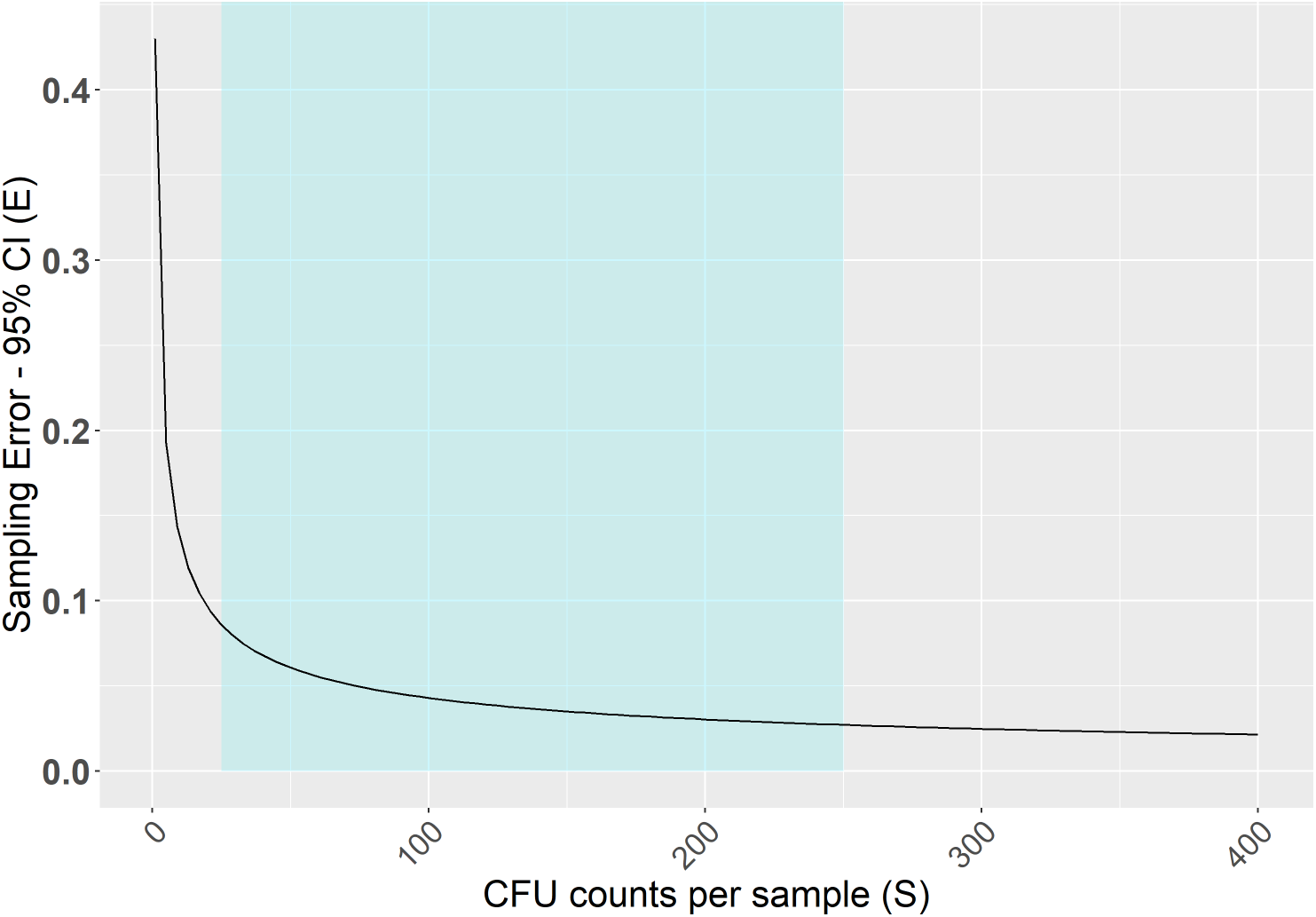
A plot of equation 3, showing the effect of CFU counts or sample size on the estimated resistance fraction’s 95% confidence intervals, or precision. Where we are under the conservative assumption that the true population resistance fraction is 0.05. The desirable range of 25-250 CFUs per sample for resistance fraction estimations is highlighted.

### Validation resistance selection environmental air samples

All isolates scored as resistant on itraconazole double-layered flamingo medium plates grew when subcultured on 4 mg/L itraconazole supplemented standard medium, while the isolates scored as resistant on the voriconazole double-layered flamingo medium plates grew when supplemented on 2 mg/L voriconazole supplemented standard medium. The two *cyp*51A gene haplotypes known to be most common in the environment, tandem repeats of either 34 (TR_34_) or 46 (TR_46_) basepairs [36], were dominant among the resistant isolates (51/58). TR_34_ was more abundant than TR_46_ (42/51) (see table S3). TR_34_ haplotypes are known to confer high-level resistance to itraconazole and often have lower-level cross-resistance to voriconazole [26, 37]. The reverse is true for TR_46_ haplotypes, which confers high-level resistance to voriconazole, and often lower-level cross-resistance to itraconazole [26, 37]. This higher abundance of TR_34_ relative to TR_46_ haplotypes in air samples is in line with what has been observed among airborne spores in the UK [18]. The ratio between the two haplotypes seems to more closely resemble that of European clinical isolates [37, 35, 38, 39] than the ratio observed in known environmental resistance hotspots sampled in the Netherlands, where TR_46_ haplotypes were more common [40]. These phenotypic and genotypic data, combined with our earlier described validation experiment, demonstrate that the described culturing method is reliable for resistance fraction estimations in environmental air samples.

### Regional baseline resistance

Figure 5 shows the estimated resistance fractions to itraconazole [A] and voriconazole [B] per sampled region. For both triazoles baseline resistance fractions ranged from 0-0.1. We found no significant regional differences for either screened triazole. Triazole resistant *A. fumigatus* is known to commonly be cross-resistant to multiple 14*α*-demethylase inhibitors (DMIs) [26]. In light of this, our findings are comparable to those of multiple European studies assessing resistance in both soil and air which reported similar resistance fractions ranging between 0 and 0.1 [41, 42, 17, 43, 18]. Given that neither air nor soil are prolific growth environments for *A. fumigatus*, we hypothesise that such values represent local baseline resistance levels present in the environment rather than resistance hotspots such as agricultural plant waste piles containing residues of triazoles.

**FIG 5.**
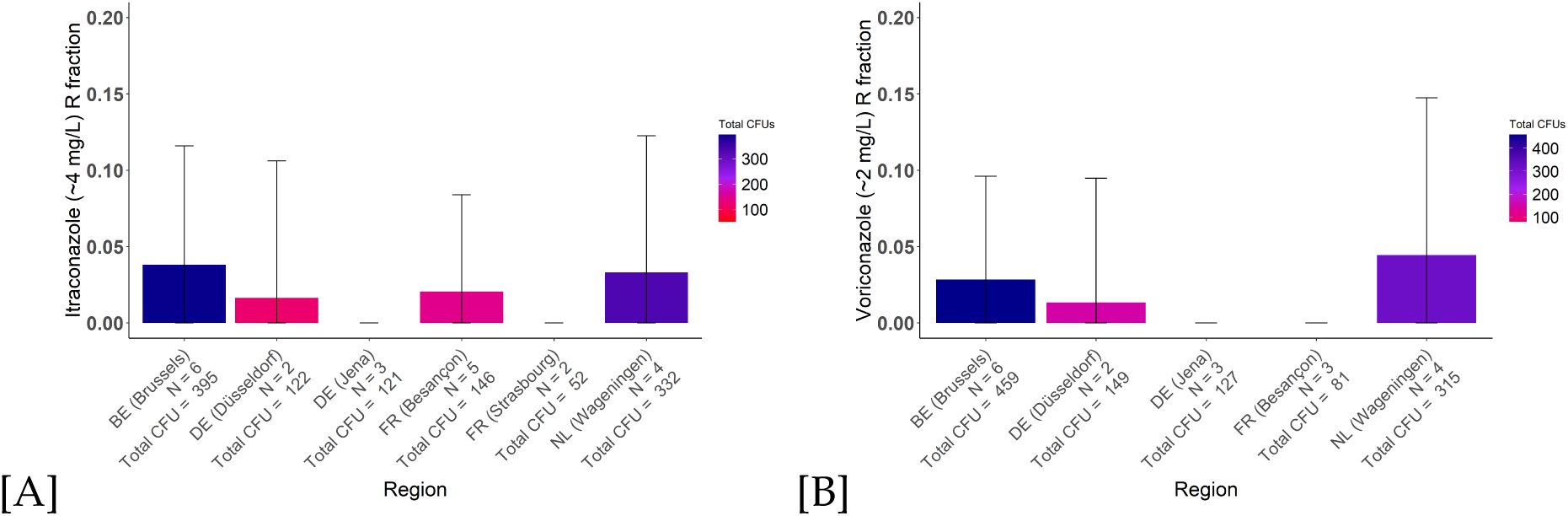
Itraconazole (*∼*4 mg/L) [A] and voriconazole (*∼*2 mg/L) [B] resistance fractions per region (mean *±* SE). Only regions where multiple samples used for resistance fraction estimations grew *≥*25 CFUs, were included.

The fact that air sampling efforts have so far yielded similar antifungal resistance fractions [17, 18] seems to indicate that spatial differences in airborne triazole resistance are absent. However, the air is not a growth environment of *A. fumigatus*, but rather the medium through which it disperses. The airborne *A. fumigatus* population represents a product of many growth environments under the influence of atmospheric mixing [18]. The effects of resistance hotspots on aerial resistance fractions may therefore be subtle or relatively local. To detect such spatial differences a method that can both provide precise point estimations of antifungal resistance and practically be used for large-scale spatial sampling is needed. Our pilot study provides proof of concept that the presented method is suitable for such large-scale air sampling efforts.

### Hotspot or coldspot?

With this new tool for large-scale environmental air sampling, we argue for a sharper definition of triazole resistance hotspots [44, 36]. That is, resistance hotspots are *A. fumigatus* growth environments in which the resistance fraction to triazole(s) increases significantly above the aerial baseline resistance. Such hotspots may be considered potential health hazards if they, in turn, significantly increase the local aerial triazole resistance fraction and the prevalence of triazole resistant *A. fumigatus* spores in their immediate surroundings above the regional baseline resistance level. Peaks in aerial antifungal resistance may be linked to known and unknown hotspots by combining spatial data on aerial resistance with spatial data on land use (type of agriculture) and economic activity.

### Outlook

Here we provide proof concept of a combination of air sampling and a culturing method that holds the potential to assess aerial resistance fractions through space. The effectiveness of the method, its relatively low cost, simplicity of use in the field and thus suitability for use by citizen scientists [45], make this method suitable for large-scale air sampling efforts. This may prove key to bridging the knowledge gap in understanding the transmission of triazole-resistant *A. fumigatus* from the environment to patients.

Numbers of airborne *A. fumigatus* appear to vary through space. Therefore, pilot studies will need to precede large-scale spatial sampling. In global regions with lower (<25) CFU yields from the traps adjustments in exposure time and sampling window will have to be made to provide higher yields and reliable resistance fraction estimations. Furthermore, due to the greater technical complexity of culturing in layered agar directly from a sticky seal, the described method is not suitable for all triazoles. These unsuitable triazoles include tebuconazole which has been used for screening in previous air sampling studies [17, 18]. These mixed results highlight the need for further validation preceding the application of other fungicides in the here-described culturing approach. Nonetheless, having an efficient and accessible air sampling and resistance screening protocol opens up extensive options for surveillance and research. The impact of known resistance hotspots [44, 40] on local aerial resistance levels can now be quantified. This is key for evaluating preventive measures where they matter most, in the air.

## MATERIALS AND METHODS

### Validation layered selective culturing

To test whether the selection for triazole-resistant colonies was effective when culturing with BIORAD^TM^ microseal seals ‘B’ (MSB1001), hereafter called ‘seals’, we selected eight *A. fumigatus* strains with known, contrasting MIC_95_ values for itraconazole and voriconazole (Fig. 1). We selected an additional set of four strains with contrasting MIC_95_ values for difenoconazole and tebuconazole for a similar test with these compounds (See Table S1). Spore suspensions were made with 0.05% saline-Tween. This solution was prepared by adding 500 µl tween-80 to 1 l 0.9% saline solution. We diluted the spore suspension such that they contained 1-40 spores per 75 µl. We cut seals into small fragments (4 cm x 6,85 cm), such that they could be placed in 9 cm Petri dishes.

Within a levelled laminal flow hood, we then placed the sticky seals inside of the Petri dishes, the sticky side facing up and exposed. 75 µl of spore suspension was pipetted in droplets across the seal to ensure colony spread. We left the droplets to dry on the seal for an hour under a laminar flow hood. We prepared Flamingo medium as described in (Zhang et al. 2021) except that, after the autoclaving step, we let the medium cool to 60 °C before adding the antimicrobial supplements and 12.5 ml of 2M sucrose as a carbon source. Subsequently, 24 ml of Flamingo medium was poured into the plate so that the seal and bottom of the plate were covered with an 8mm layer of medium. This ensured that the agar was liquid but still within the temperature tolerance range (Kozakiewicz and Smith 1994) of *A. fumigatus*. We left the agar to solidify at room temperature and incubated the plates at 48 °C for 30 hours, then kept them at 4 °C overnight, and cultured them for 8 more hours at 48 °C. Following this intermittent incubation, we marked all visible colonies and added the second layer of Flamingo medium of 12 ml (4 mm thickness) containing either no triazoles, 4 mg/L itraconazole or 4 mg/L voriconazole to plates from a set of plates for each of the reference strains. The plates were then incubated for six more days at 48 °C and the growth was scored on the agar surface.

### International Delta trap air sampling

During November 2022 a total of 70 air sampling packages were distributed or posted across Western Europe to the participants, who had no prior experience using the air sampling method. The sampling packages contained a foldable delta trap (Biogrow, Leuven, BE), a zip-lock bag, containing three sticky seals (of which the length was cut down to 11 cm by removing one of the non-sticky ends), a piece of paper to note down postal code, country and sampling start and end dates, a 30 cm piece of rope to tie up the delta trap, a strip of six poster putties, and sampling instructions. All of these materials were packaged in a 22.9 by x 33.4 cm envelope. Participants were given instructions to place the sticky seals in the delta trap and expose them for four weeks in an outdoor location near their homes. After exposure, the participants re-covered the seals and returned them via post. For a detailed description of the sampling package and the sampling instructions, including a link to the instruction video, see Supplement - Delta trap air sampling.

### Resistance screening protocol environmental air samples

To test the applicability of the layered culturing approach in environmental samples, we modified the layered culturing approach to selectively culture *A. fumigatus* from sticky seals that had been exposed to outdoor air. The volumes of Flamingo medium were adjusted to 60 ml for the initial triazole-negative flamingo medium, and 30 ml for the triazole-treatment layer. We did so to retain 8 mm and 4mm thickness, respectively, of the agar layers in larger square Petri dishes with vents (120 mm X 120 mm X 17mm) that can accommodate a sticky seal of which one non-sticky strip has been cut such that all seals are 11 cm long. Furthermore, the seals are attached to the bottom of the agar plate with Pritt^TM^ Compact Adhesive Rollers to keep the larger sticky seals flat. We adjusted the initial incubation times to 16h at 48 °C, six hours at 4 °C, and 20 hours at 48 °C. After adding the secondary triazole layer, the plates were incubated for 24 more hours at 48 °C. At this time, the total number of CFUs on the plates was counted. Finally, plates were incubated for seven more days at 48 °C after which the sporulating colonies were counted as resistant. The number of resistant colonies per seal (R) and the total number of CFUs (Tot) per seal can be used together to estimate a resistance fraction (RF) by RF = R/Tot. For a detailed day-by-day description including pictures of the environmental air sample resistance screening protocol and its rationale, see Supplement - Layered culturing protocol.

### Validation resistance screening environmental samples

To validate the selectivity of our culturing approach, we selected colonies from environmental air samples which visually sporulated with the unaided eye within nine days following the start of incubation. We selected 28 and 30 colonies from voriconazole and itraconazole plates, respectively. Colonies from voriconazole and itraconazole plates were transferred to a set of triazole-negative growth control slants, and slants containing 2 mg/L of voriconazole or 4 mg/L of itraconazole, respectively. We used 2 mg/L voriconazole to validate the resistance of colonies from the voriconazole plates rather than the 4 mg/L used in the selective layer because in earlier validation experiments the rapid (within 20 hours) suppression of growth of colonies of voriconazole-susceptible strains in the permissive layer indicated more diffusion through permissive agar layer than in the itraconazole plates. This observed greater diffusion corresponds to the 10x higher solubility in water of voriconazole than itraconazole [20, 21]. Due to this diffusion, the effective concentration in the double-layered voriconazole plates is lower than 4 mg/L, likely closer to 2 mg/L. To validate clinical resistance, the concentrations of itraconazole and voriconazole in the slants were chosen based on the EUCAST clinical breakpoints for these compounds [29]. The slants were subsequently incubated at 37 °C and scored for growth after four days. The isolates were then genotyped by harvesting spores from the growth control slants and extracting DNA using the protocol described in Fraczek et al. 2019 [46]. To detect known tandem repeat haplotypes we designed a new primer combination which allows for genotyping different TR types on agarose gel without the need for Sanger sequencing. We used this primer combination to amplify the promoter region of *cyp*51A as a resistance marker. For the PCR protocol, see Supplement - PCR protocol *cyp*51A genotyping.

### Data analysis

For the analysis of the CFU yield per seal, we used a negative binomial GLM with region as a fixed effect. Weather data was extracted from "ERA5 monthly averaged data on single levels from 1940 to present" database of the Copernicus Institute. To model weather effects on the CFU yield we included Total precipitation, 2m temperature, 10m windspeed and Downward UV radiation at the surface as fixed effects. A post hoc Tukey test was performed to test the significance of pairwise differences. For the analysis of the resistance fraction data per region, we used binomial GLMs. Statistical analysis and data visualisation were performed in R version 4.3.1 [47], using the following R packages: AER 1.2-10 [48], AICcmodavg 2.3-2 [49], cmsafops 1.3.0 [50], dplyr 1.1.2 [51], emmeans 1.8.8 [52], ggplot2 3.4.3 [53], ggpubr 0.6.0 [54], lme4 1.1-34 [55], MASS 7.3-60 [56], multcomp 1.4-25 [57], multcompview 0.1-9 [58], ncdf4 1.21 [59], rgdal 1.6-7 [60], tidyr 1.3.0 [61], rnaturalearth 0.3.4 [62], rnaturalearthdata 0.1.0 [63], raster 3.6-23 [64], sf 1.0-14 [65, 66].

## Supporting information

Detailed Delta trap air sampling protocol including how to make the package, video instructions on how to use the delta traps for air sampling.

A detailed day-by-day description of the including pictures of the environmental air sample resistance screening protocol, using double-layered agar.

PCR protocol for TR-genotyping of the cyp51a gene as a quick way to screen for resistance genotypes without the need for sequencing.

## ACKNOWLEDGMENTS

We want to thank Bart Pannebakker, Frank Becker, Ben Auxier, Eric Bastiaans, Jianhua Zhang from the Laboratory of Genetics and the other members of the Groen III One Health Consequences of Circularity team for their helpful suggestions and critical discussions. We thank Bo Briggeman for filming and editing the revised Delta trap instruction videos. We also want to thank all our collaborators and fellow scientists who contributed to the international air sampling pilot. We finally thank the BSc students who helped conceptualise and test the air sampling method during the MEE course at Wageningen University & Research during spring 2022 and spring 2023.

## DATA AVAILABILITY STATEMENT

The R code and data are made available at: https://git.wur.nl/korte058/catching_more_air.

## FUNDING

This research was funded by The Netherlands Organization for Scientific Research (NWO) under the Groen III program (GROEN.2019.002): One health consequences of circularity—What lessons to learn from the saprophytic and human pathogenic fungus *Aspergillus fumigatus*?

## CONFLICTS OF INTEREST

The authors declare no conflict of interest.

## Notes

https://git.wur.nl/korte058/catching_more_air

### Competing Interest Statement

The authors have declared no competing interest.

### Summary of Updates

Major revision of the culturing protocol, its validation, inclusion of pilot data, and more detailed user-friendly description of the sampling and culturing protocol. A near complete rewrite of the previous version.

