## Supplementary material for "Catching more air: A method to spatially quantify aerial triazole resistance in *Aspergillus fumigatus*": Detailed Delta trap air sampling protocol including how to make the package, video instructions on how to use the delta traps for air sampling.

***Aspergillus fumigatus* low-cost and easy to use air sampling protocol**

**Introduction**

A challenge when quantifying aerial resistance fractions in *Aspergillus fumigatus* through space is the need for enough airborne spores per sample.
Indeed, to estimate resistance fractions quantitatively from a practically infinite population, at least 25-250 spores (see main text) should be assayed per sample per triazole screened. This protocol aims to overcome this challenge by providing a method to collect large numbers of airborne colony-forming units (CFUs) which can directly be subjected to phenotypic antifungal resistance screening. For phenotypic resistance screening, sampling will be done in sets of three sticky seals per employed plastic delta trap. Three seals easily fit into the delta traps which shield the seals from direct sunlight (ultraviolet radiation) which is harmful to *Aspergillus fumigatus* spores, while allowing air to flow through. By culturing the *A. fumigatus* conidia that stick to the seals directly from the seals, we can assess resistance fractions and isolate resistant colonies against two different triazoles per sample (voriconazole and itraconazole). The third seal may serve for the isolation of triazole-sensitive strains and for taking *A. fumigatus* population samples. Importantly, the relative convenience of this method, such as its low cost and its suitability for distribution via standard postal services, makes it suitable for large-scale applications such as surveillance screening and point prevalence measurements involving many participants and locations.

**Preparing delta trap air sampling kits**

For each sampling kit prep the following components:

- A folded delta trap wrapped (Biogrowi *https://www.biogroei.nl/deltaval-met-bodemlijmplaten* or similar) with 3 rubber bands.
- A piece of rope (~40 cm).
- A small (~15x24cm or similar) zip-lock bag containing:
  - Three PCR Seals (BIO-RAD, Ref: MSB1001), of which one non-sticky strip has been cut such that all seals are 11 cm long. (this ensures the seals fit in 12x12cm dishes for processing)
  - 6-7 poster putties (Pritt poster buddies or similar).
  - A small sample sheet on which the following metadata can be noted down:
    - Sampling start date.
    - Sampling end date.
    - Other relevant metadata such as location or remarks as required.
  - Sampling instructions, ideally in video format (if samples are taken by third parties).

These contents may be packaged in a standard C4 size envelope, which can be readily distributed by post.

**Important**: Mark the yellow cover of the seals with the sample ID before sending out the traps. This is not just to keep track of which seals come from which delta trap/sample, but also to indicate the side that it should be facing up for the participants when placing the cover back onto the seal after exposure. If the cover is put back upside down, it will become impossible to remove and the seal will be impossible to process. Hydrophobic parchment paper used for oven baking is a common household product that may be used to replace the yellow cover if it is lost. Other types of paper or plastic cannot be used.

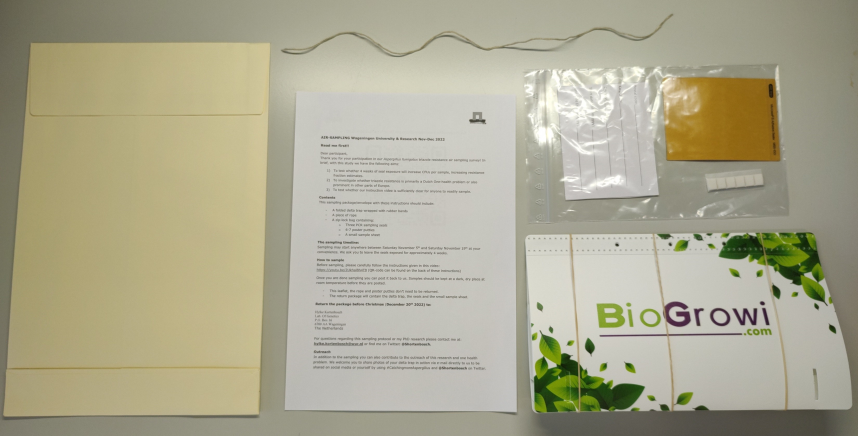

**Sample site selection**

The sampling site at which the trap is deployed should be:

- Outdoors
- Out of reach of young children or pets
- Remain undisturbed for 4 weeks after the trap is deployed
- Provide an anchor point to securely tie up the trap, for example:
  - A tree branch
  - Under a windowsill
  - A bush
  - A laundry line
  - A balcony railing

**Disclaimer regarding the use of the traps indoors or at composters**

The traps are intended for outdoor use and have only been validated to yield sufficient CFUs for resistance fraction estimations in outdoor environments after four weeks of exposure.

Indoors, the seals (without the delta trap) will generally capture at least 2-3x fewer CFUs over four weeks (data not shown). This may depend on how modern the building is and especially its ventilation/HEPA air filtration systems. Extending the exposure time beyond four weeks may mitigate the lower indoor CFU yield but this requires further validation. Use this air sampling method indoors at your discretion.

Deployment of the traps next to an *A. fumigatus* growth environment that is actively disturbed, such as when plant waste heaps are turned during composting processes, should be done with caution. This may cause too much growth on the seals with CFUs, preventing the accurate counting of CFUs and calculation of a resistance fraction from them becomes difficult. Depending on the aim of the study, for sampling such bursts of airborne *A. fumigatus,* active air sampling devices may be more practical.

**Link to playlist of video instructions on use of the delta traps air sampling method:**

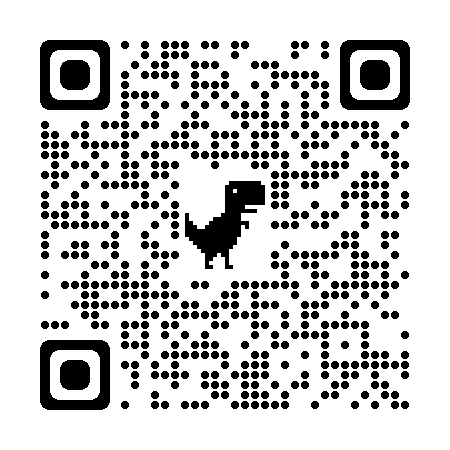

**https://www.youtube.com/playlist?list=PLWgOstKjDQFQw5RTFN7C4dQnFCDaLWDnu**

**#Sampling instructions as sent along with packages international pilot study# #including original instruction video#**

**Air sampling international pilot Wageningen University & Research Nov-Dec 2022**
**Read me first!!**
Dear participant,
Thank you for your participation in our *Aspergillus fumigatus* triazole resistance air sampling survey! In brief, with this study, we have the following aims:

1. To test whether 4 weeks of seal exposure will increase CFUs per sample, increasing resistance fraction estimates.
2. To investigate whether triazole resistance is primarily a Dutch One-health problem or also prominent in other parts of Europe.
3. To test whether our instruction video is sufficiently clear for anyone to readily sample.

**Contents**
This sampling package/envelope with these instructions should include:

- A folded delta trap wrapped with rubber bands
- A piece of rope
- A zip-lock bag containing:
  - Three PCR sampling seals
  - 6-7 poster putties
  - A small sample sheet

**The sampling timeline:**
Sampling may start anywhere between Saturday, November 5^th^ and Saturday, November 19^th^ at your convenience. We ask you to leave the seals exposed for approximately 4 weeks.
**How to sample**
Before sampling, please carefully follow the instructions given in this video: <https://youtu.be/2Ukhal8h4T8> (The QR code can be found on the back of these instructions)
Once you are done sampling you can post it back to us. Samples should be kept in a dark, dry place at room temperature before they are posted.

- This leaflet, the rope and poster putties don’t need to be returned.
- The return package should contain the delta trap, the seals and the small sample sheet.

**Return the package before Christmas** (**December 20^th^ 2022) to:**
Hylke Kortenbosch
Lab. Of Genetics
P.O. Box 16
6700 AA Wageningen
The Netherlands
For questions regarding this sampling protocol or my PhD research please contact me at:  

**For the sampling instructions please see the other side of this sheet**

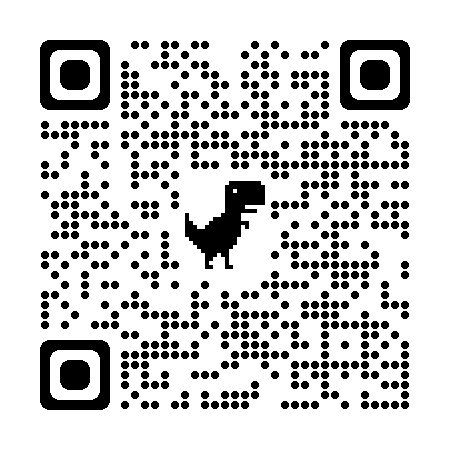
