## Supplementary material for "Catching more air: A method to spatially quantify aerial triazole resistance in *Aspergillus fumigatus*": A detailed day-by-day description of the including pictures of the environmental air sample resistance screening protocol, using double-layered agar.

### Supplement – Layered culturing protocol

**Detailed description of layered culturing protocol for *A. fumigatus* antifungal resistance screening of outdoor air samples**

#### Introduction

This protocol is used to analyse delta-trap air samples of the airborne mould *Aspergillus fumigatus* for triazole resistance screening and further downstream analysis of individual colony-forming units (CFUs). A major issue with accurately estimating antimicrobial resistance fractions using culturing methods is the need for growth-control plates to represent the total number of CFUs per sample on a non-selective agar plate compared to an antimicrobial selective agar plate, in our case, triazole compounds. This leads to two estimates and thus two sampling errors, one for the total number of CFUs per a non-selective agar plate and one for the number of resistant CFUs on the antimicrobial selective agar plate. By utilising a layered culturing approach, we integrate the growth control non-selective and antimicrobial selective agar plates into a single plate per air sample seal, eliminating the sampling errors as indicated above, and increasing the accuracy of our resistance fraction estimation.

The aim of this protocol is to selectively culture for *A. fumigatus* such that both the total number of CFUs as well as the respective number of voriconazole (MIC > 1 mg/L), itraconazole (MIC > 2 mg/L) resistant CFUs, may be counted on two of the three available seals sampled per delta trap. The 3rd seal may be used as an additional azole-negative control plate to grow individual CFUs (including triazole-sensitive colonies) from the total population for further downstream processing. The EUCAST thresholds for voriconazole (>1 mg/L) and itraconazole (>2 mg/L) resistance were chosen as reference points when choosing the used triazole concentrations. Incubation and culturing times were chosen assuming 8-hour workdays and a weekend on Saturday/Sunday.

#### Preparation: The Flamingo medium

To ensure all viable spores on the seal germinate, an initial layer (without azoles) of Flamingo medium is added. To selectively culture *Aspergillus fumigatus* we use Flamingo medium as described in Zhang et al 2021. The minimal medium for aspergilli forms the basis of the Flamingo medium and by supplementation with rose bengal and dichloran, non-*Aspergillus fumigatus* fungal growth is inhibited.

Components Minimal Medium for *Aspergilli* (1L)

| **Component** | **Amount** | **Units** |
|---|---|---|
| $NaNO_3$ | 6.0 | gram |
| $KH_2PO_4$ | 1.5 | gram |
| $MgSO_4.7H_2O$ | 0.5 | gram |
| KCl | 0.5 | gram |
| $FeSO_4$ | 1 | mg |
| $ZnSO_4$ | 1 | mg |
| $CuSO_4$ | 1 | mg |
| $MnCl_2.4H_2O$ | 1 | mg |
| Agar | 15 | gram |
| Sucrose | 8.6 | gram |
| Demin | 1 | litre |

- Prepare a 10 mg/ml master mix of $FeSO_4$, $CuSO_4$, $ZnSO_4$ and $MnCl_2.4H_2O$ in demineralised water and add 1 ml master mix to medium (Given the small quantities, making 30 ml for long-term use is recommended)
- Weigh the proper amount of $NaNO_3$, $KH_2PO_4$, $MgSO_4$ .$7H_2O$, KCl and sucrose.
- Dissolve in a final volume of 1 litre demi water.
- Adjust pH to 5.8
- Add the agar
- Autoclave at 120 °C for 20 min.
- MM for *Aspergilli* keeps well at room temperature for months when stored appropriately and may be prepared well in advance of processing the air samples.

Note: When preparing the medium in larger batches it is recommended to weigh and mix all the salts in a large pot, and then adjust the pH. This mixture can then be added to 1-litre bottles to which the agar and sucrose have already been added. This way, the pH must be adjusted only once and only the agar and sucrose need to be weighed separately per 1-litre bottle.

**Adding antimicrobial supplements to make Flamingo medium out of MM**

- Add 10 ml Dichloran stock solution and 5 ml Rose Bengal stock solution (final concentration of 10 mg/l Dichloran solution and 25 mg/l Rose Bengal, respectively) to 1 L MM wearing gloves and mix gently by inverting the bottle. **Be careful when handling the Dichloran solution, the acetone is volatile and highly flammable! Wear gloves when handling the Rose Bengal as it is a powerful staining agent.**

<u>Preparation of a 100 ml (5 mg/ml) Rose Bengal stock</u>:

- Weigh 500 mg Rose Bengal.
- Dissolve 500 mg Rose Bengal in 30 ml 95% ethanol and add demineralised water up to a total volume of 100 ml. Mix gently until fully dissolved.
- Store at 4 ºC in a bottle covered with aluminium foil to limit photochemical degradation.

<u>Preparation of a 250 ml (1 mg/ml) Dichloran stock</u>:

- Wearing gloves, weigh 250 mg Dichloran (safety cabinet W2.Ha.055).
- dissolve 250 mg Dichloran in 250 ml 95% acetone and mix carefully until fully dissolved.
- Store at 4 °C in a bottle covered with aluminium foil to avoid photochemical degradation.

Flamingo is further supplemented with the antibiotics chloramphenicol and streptomycin to suppress any bacterial background growth.

- Add 500 µl 100 mg/ml chloramphenicol (in 96% ethanol) to 1 litre Flamingo (final concentration 50 mg/l).
- Add 250 µl 200 mg/ml streptomycin (in water) to 1 litre Flamingo (final concentration 50 mg/l).
- Antibiotic stock solutions should be stored in a -20 °C Freezer.

When pouring the flamingo medium over the seals it should have been cooled down to 60 °C as higher temperatures may kill off the *A. fumigatus* spores present on the seal, and lower temperatures will have the agar solidify too fast. It is therefore recommended to melt the minimal medium and add the additives to it one day in advance of pouring the flamingo medium and leaving it in a 60 °C stove to cool overnight.

Note: Because the Dichloran is dissolved in acetone, which has a low boiling point, it should not be added to the minimal medium when it is still close to 100 °C. First, let the medium cool (not solidify) for two hours in a 60 ºC stove or otherwise cool it down before adding the dichloran and other additives.

#### Day-by-day culturing protocol

##### Day 0: Placing seal into Petri dishes + add first layer of Flamingo medium

**Preparation**

- Disinfect your workbench.
- Take your exposed seals.
- Take square Petri dishes (Greiner Bio-one, Ref: 688102) (1 per seal to be processed).
- Take Pritt adhesive roller or similar.
- Clean hands with 70% ethanol before and after handling each set of seals.

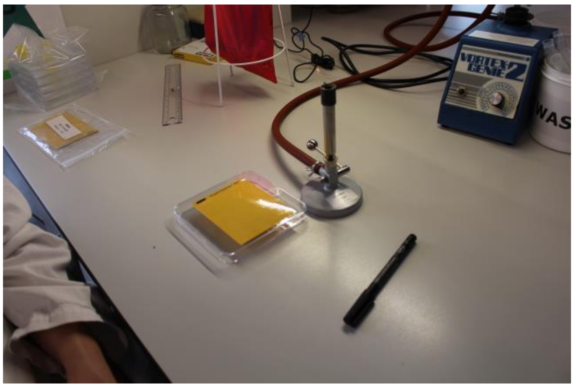


Work next to a flame or in a biosafety cabinet to avoid contamination. Ensure that the workbench is levelled, because an equal thickness of the agar layers is crucial for reliable resistance screening.

**Placing the seals into the Petri dishes**

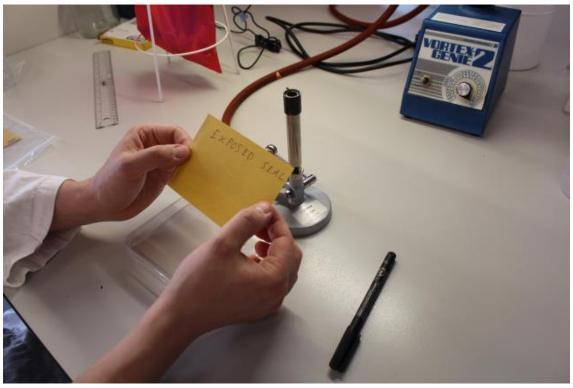


Using the Pritt adhesive roller, roll two ~10 cm lines of adhesive in parallel along the long sides of the non-sticky back of the seal. **Not its yellow cover!!** **Note**: Double-sided scotch tape also works for sticking the seals into the plates when applied similarly to the adhesive roller but is overall less practical to work with.

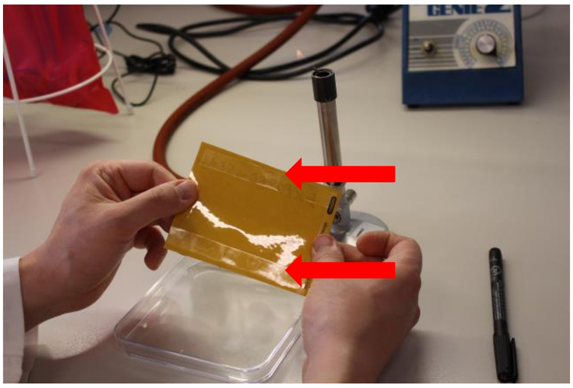

*Ad adhesive to the reflective non-sticky side of the seal (see red arrows).*

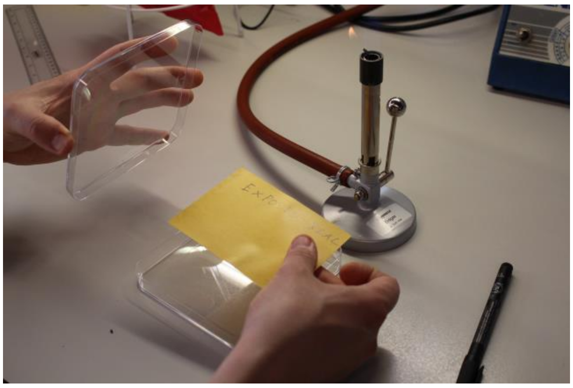

*Now place the seal on the bottom of a square Petri dish.*

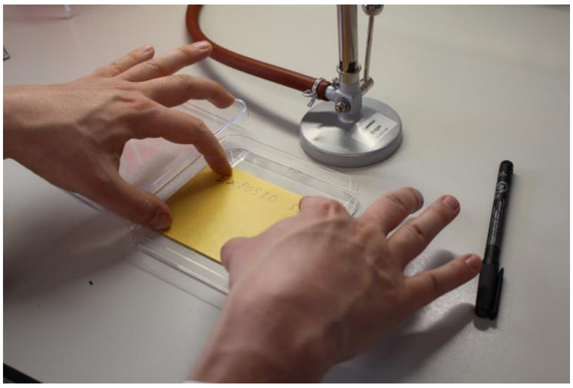

*Make sure the seal is stuck flat to the bottom of the plate by pressing gently on the yellow cover along the edges where the adhesive was applied.*

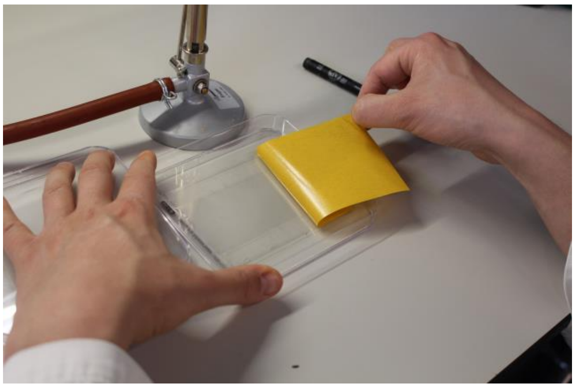

*Gently remove the yellow cover from the seal by pulling it sideways rather than upwards.*

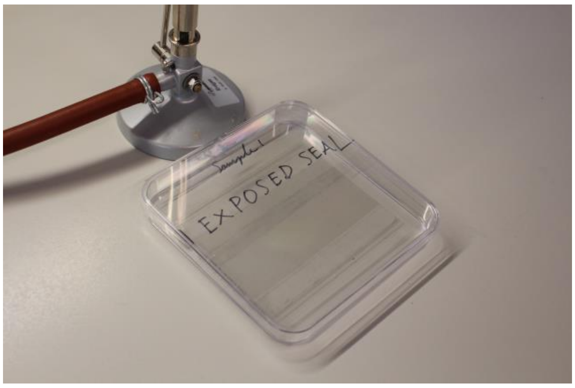


*Close and label the plate and repeat for all sticky seals that need to be processed.*

**Adding the permissive layer of flamingo medium**

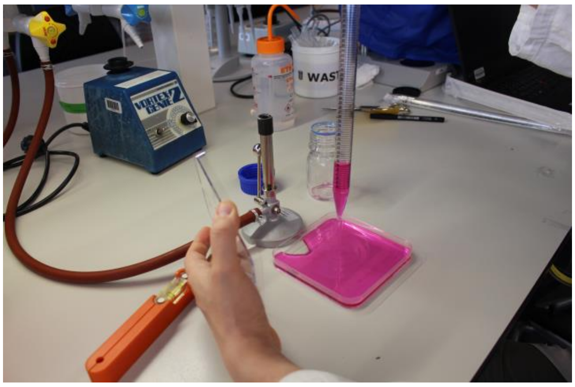

Pour a 60 ml (8 mm thick) layer of azole-negative flamingo medium over the seal, starting at its centre and leave it to fill out the plate. It should do this by itself if the workbench is levelled and the medium is at 60 °C. At this stage, it is critical to have a levelled workbench, be it a workbench or biosafety cabinet, to ensure an even thickness of the agar over the seal. For a convenient workflow pouring the plates can best be done in the late afternoon.

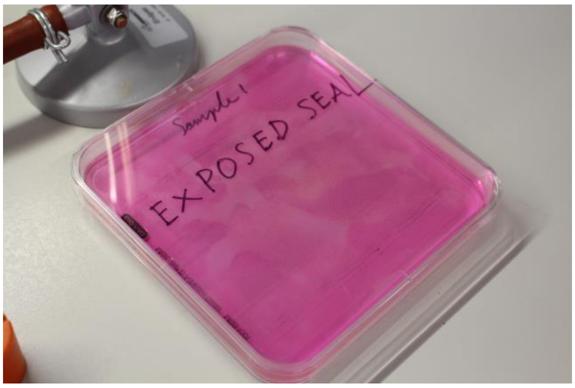


Leave the agar to solidify at room temperature for 30-45 minutes on a levelled surface, then incubate for 16 hours (overnight) at 48 °C. Starting the incubation at 5 pm works well with this time window.

#### Day 1: Changing incubation temperatures

After culturing the plates overnight (16 hours) remove the plates from the 48 °C incubator and place them at 4 °C for 6 hours, and subsequently incubate them for another 21 hours at 48 °C. If plates were originally placed in the incubator at 5 pm, they should be placed at 4 °C at 9 am, placed back in the incubator at 3 pm, and subsequently incubated till noon
Note: Placing the plates at room temperature for 15 minutes when moving the plates between 48 °C and 4 °C will reduce the formation of condensation on them.

Alternatively, the plates may be cultured for a continuous 35 hours at 48 °C. This will remove the need for moving the plates to and from a fridge, but this time window does not readily fit into regular working hours.

**Rationale culturing times**
These specific culture times are required to, first, ensure that most CFUs have started forming within the agar before the second triazole-containing layer is added. Second, at this stage, the colonies will not have breached the agar surface yet when the second layer is added, which might smear (resistant) spores across the agar surface.

Not adding the triazole layer too early is especially important before adding the voriconazole-containing medium. This compound has a relatively high water-solubility for a triazole and will diffuse throughout the plate in about a day, preventing all growth in the bottom layer if added from the start. This would prevent assessment of the total CFU count on the seal, and thus accurate estimation of the voriconazole resistance fraction.

#### Day 2: Adding the second, triazole-containing layer

After the initial incubation period in the triazole-free layer, a second layer of 30 ml Flamingo medium is added at 60 °C. The medium can contain a different triazole for each of the two or three seals of each delta trap. The two validated triazoles are Voriconazole (4 mg/L), and Itraconazole (4 mg/L). The triazoles are added to the Flamingo medium from DMSO stocks and stored at -80 °C. Add 1 ml of a 4 mg/ml stock to 1 l of Flamingo medium to get the desired triazole concentration.

Again, work under sterile and levelled conditions:
Pour the medium onto the centre of the plate, if the workspace is levelled and the medium at 60 °C, it should spread over the bottom of the plate on its own.
Leave to solidify for 30-45 minutes at room temperature, note that smaller stacks will solidify faster than larger ones. Stacks of three plates appear a good compromise between time and space efficiency.  If working at the advised times, adding the second layer should be start at around noon.

Incubate for 16-20 hours at 48 °C - the effect of the applied triazole is still minimal at the bottom of the permissive layer during this time so the final colonies may still appear.

#### Day 3: counting CFU totals per seal

On day 3, the CFUs on the seal should have started visibly growing.

The colonies will still be small at this stage, many having a diameter no greater than ~ 3mm. Therefore, we recommend the use of both a strong backlight and front light (or sunlight) in combination with a dark background to count and mark the colonies with a marker on the plate while counting, this will be the total CFU count.

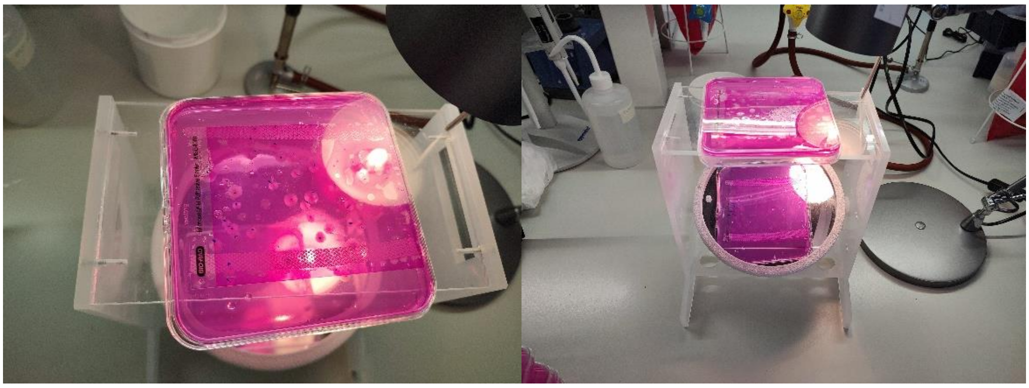

For counting the colonies on the bottom of the plates, the plates may be flipped or tilted over as the colonies will not be sporulating yet, so spore scattering is not an issue. On the control plates without the second layer, some colonies may have breached the surface after 3 days. These plates should thus be handled with extra care when counting CFUs. Especially for the control plates, counting should be started at 9 am or otherwise at the start of the workday so that colonies will still show minimal sporulation and the risk of spore scattering is limited when flipping the plates to count the colonies. Timing is less critical for the selective triazole plate as the agar on those is both thick and contains a triazole. Nonetheless, counting in the morning is advised.

Flamingo medium is selective for *A. fumigatus* but several types of mucor colonies may still grow. These are, however, clearly distinguishable from *A. fumigatus* colony morphology, also when colonies are still small. Typically, mucor colonies will be larger and less dense than the *A. fumigatus* colonies on day 3. See the examples below in which the *A. fumigatus* colonies are marked with black dots and the mucor with blue circles. The small black dots mark scored *A. Fumigatus* colonies.

**Note**: if mucor is present on a seal, it will have breached the agar surface by day 3. To prevent smearing it across the plate, it is recommended to, if possible, use plates with mucor as itraconazole plates, rather than voriconazole plates. Mucor species have little to no inhibition by voriconazole, while they are more inhibited by itraconazole, so cause fewer issues on the latter type of plates. This is not always feasible when processing large batches of samples but is something to consider for single samples or small batches.

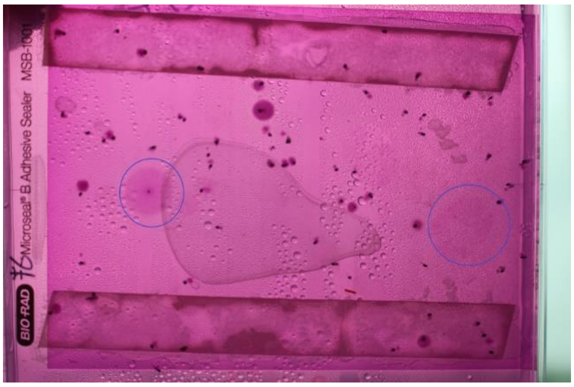


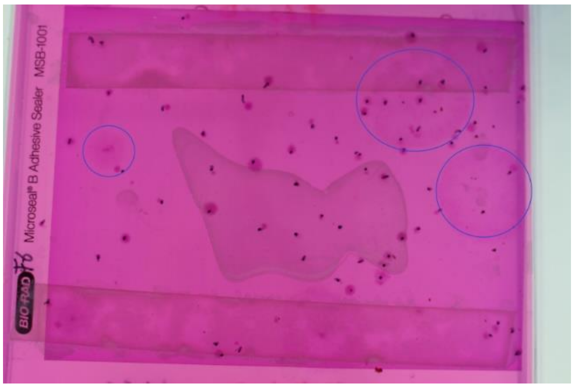


Continue the incubation of the plates at 48 °C.

#### Day 5 - 6: Remove growth control plates from the incubator

By day 5-6, any plates that did not have a second selective triazole layer added should now have sporulating colonies which may be used to collect a population sample or otherwise be analysed. On day 5 there will be less overlap between colonies which will make single colony isolation easier, while on day 6 they might be too big and no longer individually spaced colonies.

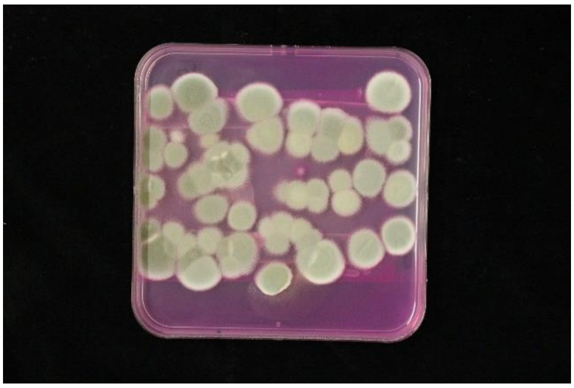

An example of a growth control plate at day 6 of incubation that should not be incubated further but stored at 4 degrees.

#### Day 9: Count the number of resistant colonies per seal

By day 9, count individual colonies on the selective triazole plates that have breached the top layer, and are visually sporulating, as resistant. Validation experiments have shown that non-sporulating colonies are not consistently resistant to 2 mg/L itraconazole or voriconazole when transferred to slants and should conservatively not be counted as resistant colonies.

The resistance fraction can be calculated per seal: # of resistant colonies / total # of colonies (of the same plate on day 3) = resistance fraction.

If colonies were marked clearly on day 3 when counting the totals, overlapping colonies would typically still be distinguishable individually.

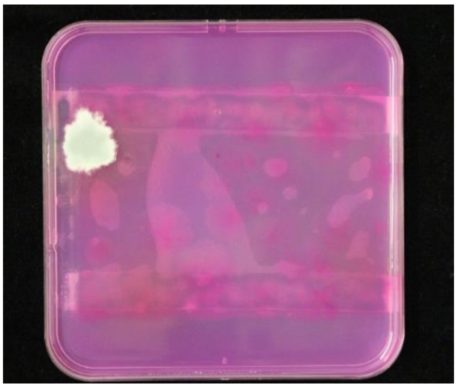

An example of an itraconazole plate with a single resistant colony

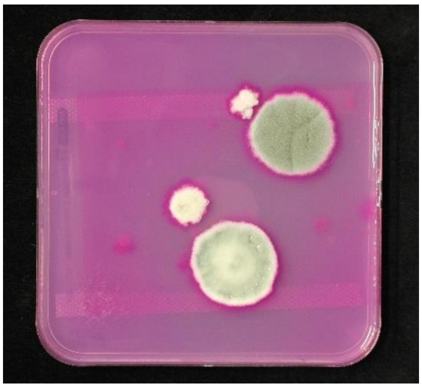

An example of a voriconazole plate with two resistant colonies, the two smaller non-sporulating colonies on the top left of the two larger colonies are not counted as resistant.

**Example of output data in long format**

For these samples, two seals were screened for resistance to itraconazole (ITR) and voriconazole (VOR), respectively. The third seal was used as a growth control.

| Sample | Total ITR | RES ITR | RF ITR | Total VOR | RES VOR | RF VOR | Total control |
|---|---|---|---|---|---|---|---|
| A1 | 41 | 2 | 0.04878 | 56 | 3 | 0.053571 | 62 |
| A2 | 91 | 3 | 0.032967 | 82 | 1 | 0.012195 | 95 |
| A3 | 224 | 6 | 0.026786 | 200 | 1 | 0.005 | 220 |
| A4 | 82 | 5 | 0.060976 | 122 | 1 | 0.008197 | 84 |
