## Supplementary material for "Catching more air: A method to spatially quantify aerial triazole resistance in *Aspergillus fumigatus*": PCR protocol for TR-genotyping of the cyp51a gene as a quick way to screen for resistance genotypes without the need for sequencing.

**Supplement - PCR protocol genotyping *A. fumigatus* *cyp51A* TR-types**

Primer sequences:

| Primer | Name | Sequence (5'-3') |
| --- | --- | --- |
| Forward | TRC51_F1 | AATCGCAGCACCACTTCAGA |
| Reverse | 365.1 | CCATAGCATCGGCACCAT |

Master mix:

| Component | Volume (uL) (1x) |
| --- | --- |
| 5xGoTaq mix | 5 |
| dNTP 5mM | 0.5 |
| Primer F (TR51-F1) | 0.5 |
| Primer R (365.1) | 0.5 |
| Template | 3 |
| GoTaq | 0.1 |
| H2O | 15.4 |
| Sum volume | 25 |

PCR programme:

| 1x | 2 min | 95°C |
| --- | --- | --- |
| 35X | 20 sec | 95°C |
|  | 20 sec | 58°C |
|  | 1 min | 72°C |
| 1x | 5 min | 72°C |
| 1x | Hold | 12°C |

The PCR amplicon fragment length is in the range of ~360-410 bp for the common *cyp*51A Tandem Repeat genotypes: WT, TR34 and TR46. This enables genotyping of these common environmental genotypes on an agarose gel without Sanger sequencing. The PCR product was run in 2% agarose gels for 120 minutes (2 hours) at 130 V for clear separation of the TR34 and TR46 bands (see picture below).

Example of gel ran with described PCR protocol and gel electrophoresis:

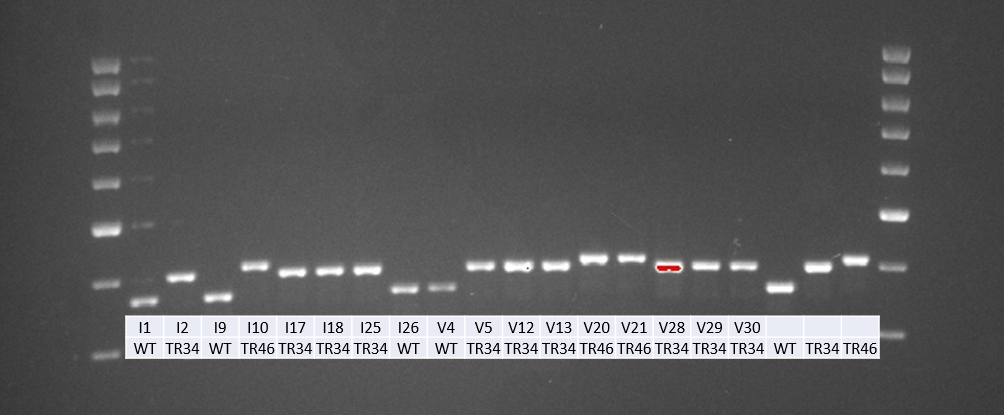
