## Supplementary Tables S1 to S4 for "Catching more air: A method to spatially quantify aerial triazole resistance in *Aspergillus fumigatus*"

**Supplemental tables**

Table S1: Growth of characterised reference strains though a top layer which either included no triazole (0), 4 mg/L Difenoconazole (DIF 4) or 6 mg/L Tebuconazole (TEB 6) MIC95 values of the strains to the respective triazole are given. There is some inhibition by the triazoles of the sensitive strains, but this is quantitative rather than qualitative.

| **Strain** | **Genotype** | **MIC DIF** | **MIC TEB** | **0** | **DIF 4** | **TEB 6** |
| --- | --- | --- | --- | --- | --- | --- |
| **55C1** | **Wt** | **0,5** | **0,5** | **+**  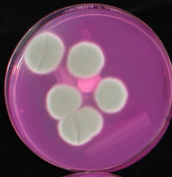 | **+**  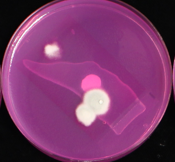 | **+**  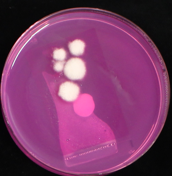 |
| **78C1** | **Wt** | **0,25** | **0,5** | **+**  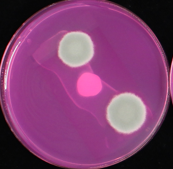 | **+**  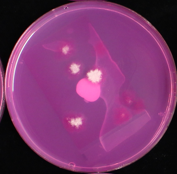 | **+**  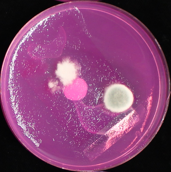 |
| **11A6** | **TR34** | **8** | **4** | **+**  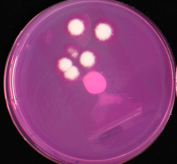 | **+**  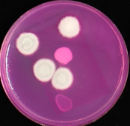 | **+**  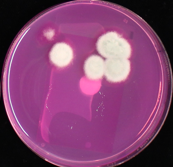 |
| **25C20** | **TR46** | **2** | **>16** | **+**  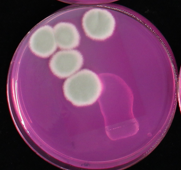 | **+**  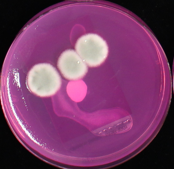 | **+**  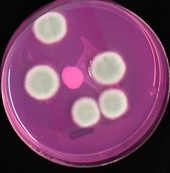 |

Table S2: Growth of characterised reference strains when plated on the agar surface with glass beads. Treatments included are: no triazoles (0), 4 mg/L Difenoconazole (DIF 4) or 6 mg/L Tebuconazole (TEB 6). MIC values of the strains to the respective triazole are given. Growth inhibition is in line with MIC values.

| **Strain** | **Genotype** | **MIC DIF** | **MIC TEB** | **0** | **DIF 4** | **TEB 6** |
| --- | --- | --- | --- | --- | --- | --- |
| **55C1** | **Wt** | **0,5** | **0,5** | **+** | **-** | **-** |
| **78C1** | **Wt** | **0,25** | **0,5** | **+** | **-** | **-** |
| **11A6** | **TR34** | **8** | **4** | **+** | **+** | **+** |
| **25C20** | **TR46** | **2** | **>16** | **+** | **+** | **+** |

Table S3: Output of the Negative binomial model used to test for an effect of region on CFU yield per seal. Output indicates a significant effect of region on the CFU yield. Post-Hoc results are visualised in Fig

|  | **Estimate** | **Std.Error** | **z value** | **Pr(>\|z\|)** |
| --- | --- | --- | --- | --- |
| Intercept | 4.30856 | 0.16695 | 25.808 | <2e-16 |
| **DE (Düsseldorf)** | -0.06723 | 0.3346 | -0.201 | 0.84075 |
| **DE (Jena)** | -0.76038 | 0.27113 | -2.805 | 0.00504 |
| **DK (Copenhagen)** | -1.41819 | 0.28321 | -5.008 | 5.51E-07 |
| **ES (Madrid)** | -1.82365 | 0.29521 | -6.177 | 6.51E-10 |
| **FI (Tampere)** | -1.5052 | 0.36751 | -4.096 | 4.21E-05 |
| **FR (Besançon)** | -1.20997 | 0.24702 | -4.898 | 9.67E-07 |
| **FR (Nantes)** | -1.10989 | 0.35379 | -3.137 | 0.00171 |
| **FR (Strasbourg)** | -1.41819 | 0.31249 | -4.538 | 5.67E-06 |
| **FR (Verton)** | -1.70587 | 0.37656 | -4.53 | 5.89E-06 |
| **NL (Wageningen)** | 0.05772 | 0.26361 | 0.219 | 0.82669 |

Table S4: Summary table showing the TR34 and TR46 cyp51A genotypes of the isolates grown on the itraconazole (ITR) and voriconazole (VOR) plates of the environmental samples taken during the international air sampling pilot.

| **Genotype** | **ITRA** | **VOR** | **Total** |
| --- | --- | --- | --- |
| **TR34** | **19** | **23** | **42** |
| **TR46** | **5** | **4** | **9** |
| **WT** | **6** | **1** | **7** |
| **Total** | **30** | **28** | **58** |
